# Feeding and reproduction of a tropical coastal copepod across warming and copper gradients

**DOI:** 10.64898/2026.03.09.710611

**Authors:** Ngoc-Anh Vu, Minh-Hoang Le, Tuan-Anh Hoang Lu, Hoang Viet Luu, Nam X. Doan, Kiem N. Truong, Khuong V. Dinh

## Abstract

Tropical coastal ecosystems in Southeast Asia are facing rapid warming and increasing pollution. Shallow coastal waters now frequently exceed 34 °C, potentially pushing tropical ectotherms beyond their thermal optimum while they are simultaneously exposed to copper (Cu) contamination, especially from aquaculture and shipping activities. However, how warming alters Cu toxicity in dominant tropical zooplankton remains poorly understood. We examined the effects of Cu (0, 10, 20, 30 and 40 µg L^−1^) and temperature (26, 29, 32 and 35 °C) across realistic gradients on the calanoid copepod *Pseudodiaptomus annandalei*, a dominant grazer of coastal plankton communities. Adult survival, cumulative faecal pellet production (as a proxy for energy intake), and cumulative nauplii production were quantified over seven days. No significant effects of temperature or Cu on adult survival were detected, likely reflecting variability among wild-collected individuals. In contrast, temperature was the main driver of feeding and reproductive performance, which peaked at 32 °C and declined at 35 °C. Cu exposure alone had no significant effects at 26 - 35 °C due to high variability in responses. At 32 °C, cumulative feeding and reproductive responses did not statistically differ among copper concentrations, whereas variability increased at both lower and higher temperatures. At 35 °C, Cu effects were non-linear, with nauplii production reduced at 30 µg Cu L^−1^ but highest at 20 µg Cu L^−1^, while faecal pellet production showed treatment-specific reductions, particularly in non-exposed individuals at 26 °C and in high Cu treatments at 29 and 35 °C. These findings indicate that warming can modify contaminant effects in tropical zooplankton and highlight the importance of incorporating realistic thermal regimes and natural population variability into ecological risk assessments under climate change.

## 1. Introduction

Tropical coastal ecosystems are increasingly exposed to multiple, overlapping anthropogenic stressors (Halpern et al., 2025, Barlow et al., 2018, Wang et al., 2013). Rapid expansion of aquaculture, urbanization, and industrialization in Southeast Asia has resulted in substantial pollutant inputs to estuarine and nearshore environments (Crain et al., 2008, Halpern et al., 2025). Among these pollutants, copper (Cu) is of particular concern due to its widespread use in antifouling paints, and in aquaculture where Cu compounds are applied as algaecides to control algal growth, and as disinfectants (U.S. Environmental Protection Agency, 2009, Ebenezer et al., 2014, Marcussen et al., 2014, Thornton and Rast, 1997). Although Cu rapidly adsorbs to particles and sediments, recurrent inputs from aquaculture and shipping activities may result in chronic exposure for planktonic organisms, where it can impair survival, feeding activity, and reproductive output in copepods and other zooplankton (Kwok et al., 2008, Dinh et al., 2020). Tropical marine ecosystems are also increasingly affected by climate warming and extreme temperature events (Lee et al., 2023, Smale et al., 2019, Gruber et al., 2021, Yao and Wang, 2021). Marine heatwaves (MHWs) have become more frequent, longer-lasting, and more intense over the past decades (Frölicher et al., 2018, Smale et al., 2019). In shallow coastal waters of Southeast Asia, temperatures of 34 °C or higher are frequently recorded during warm seasons (Doan et al., 2018). Tropical marine ectotherms typically live close to their upper thermal limits (Tewksbury et al., 2008, Deutsch et al., 2008). For example, the optimal thermal window of the calanoid *Pseudodiaptomus annandalei*, based on the respiration rate, is 26-32 °C (Lehette et al., 2016). Consequently, even small increases above their upper thermal optimum may result in disproportionate declines in survival, growth, feeding and reproduction (Nguyen et al., 2011, Doan et al., 2019). For example, *P. annandalei* shows an increased feeding rate at temperature 25-30 °C while feeding declines at temperatures 34-35 °C (Doan et al., 2018, 2019).

Importantly, warming and pollutants can interact in non-additive manners, resulting in an intensifying or alleviating combined effects (Crain et al., 2008, Dinh et al., 2022). Understanding how warming interacts with contaminants has become a major challenge in aquatic ecology and ecotoxicology in the past two decades (Sokolova and Lannig, 2008, Noyes et al., 2009, Jackson et al., 2016, Dinh et al., 2022). Most experimental studies on warming-pollutant interactions have focused on temperate species, and limited stressor levels, which do not allow for testing for the non-linear ecological responses (Dinh et al., 2022, Orr et al., 2024), even ecological surprises, unexpected or non-linear outcomes, even mass mortality caused by synergistic interaction of multiple stressors (Darling and Côté, 2008, Garrabou et al., 2009), especially at supra-optimal thermal conditions (Dinh et al., 2022, Orr et al., 2024). Several studies have examined temperature-dependent changes in metal toxicity across multiple temperatures (e.g., Bao et al., 2008, Li et al., 2014), these have primarily focused on changes in overall sensitivity rather than evaluating responses across combined gradients of temperature and contaminant levels. More broadly, the interaction outcomes of multiple stressors can vary across thermal gradients, with effects shifting from additive near optimal temperatures to more pronounced (e.g., synergistic) effects at thermal extremes (see e.g., Verheyen and Stoks, 2023). In addition, most laboratory experiments rely on highly standardized cohorts of organisms, which may not fully capture the demographic variability typical of natural populations (Rohr et al., 2016). Differences in age, reproductive stage, and physiological condition can influence organismal responses to environmental stressors, potentially altering the magnitude or detectability of contaminant effects under realistic ecological conditions (Muyssen and Janssen, 2007, Dinh et al., 2022, Schmid et al., 2023, Rohr et al., 2016). This may complicate extrapolation from laboratory assays to natural populations (Rohr et al., 2016, Dinh et al., 2022).

Despite growing recognition of temperature-dependent toxicity, few studies have evaluated contaminant effects across a realistic thermal gradient (but see e.g., Bao et al., 2008, Li et al., 2014). This gap is particularly relevant for short-lived planktonic organisms such as copepods, which form the foundation of coastal food webs (Chew et al., 2012). Calanoid copepods of the genus *Pseudodiaptomus* are dominant components of tropical plankton communities and play a critical role in transferring primary production to higher trophic levels (Truong et al., 2022, Truong et al., 2014, Chew et al., 2012). Because of their short generation time (5–12 days depending on temperature; Doan et al., 2019, Nguyen et al., 2020b), changes in survival or reproduction can rapidly alter population dynamics and ecosystem functioning.

In this study, we investigated the sensitivity of *Pseudodiaptomus annandalei* to a gradient of temperatures (26 - 35 °C) and copper concentrations (0 - 40 µg L^−1^) over seven days. By quantifying adult survival, cumulative faecal pellet production (as a proxy for food intake and energy assimilation, Isla et al., 2008), and cumulative nauplii production, we aimed to evaluate how warming and Cu exposure affect energy allocation on survival and reproduction, two of the most critical parameters for population growth. Specifically, we addressed three hypotheses (H1-H3): **H1)** Temperature has a non-linear effect on performance with feeding activity and reproductive output *P. annandalei* increasing up the thermal optimum (approximately 32 °C, Lehette et al., 2016), then declining at higher temperatures, with stronger effects on reproduction than on survival. **H2)** Exposure to elevated Cu concentrations reduces survival, feeding, and reproductive performance of *P. annandalei*. **H3)** Combined effects of temperature and Cu are non-additive and vary across a thermal gradient, particularly beyond the upper thermal optimum. By linking feeding activity and reproduction within an energy allocation framework, this study provides insights into how warming and pollution may interact in tropical coastal ecosystems.

## 2. Materials and methods

### 2.1 Pseudodiaptomus annandalei

*Pseudodiaptomus annandalei* is a dominant calanoid copepod occurring along the central coast of Vietnam and Southeast Asian regions (Truong et al., 2022, Truong et al., 2014, Chew et al., 2012). Adult males and females of *P. annandalei* were collected from an aquaculture pond in the Cam Ranh Centre for Tropical Marine Research and Aquaculture in the summer, using a zooplankton net (mesh size = 200 μm). The sampling site was located at 11°49’26.3”B, 109°07’23.9”E. At the time of sampling, the water temperature and salinity of the pond were 26 °C and 29 PSU, respectively. As the copepods were collected from a natural population, individuals likely varied in age, reproductive stage, and physiological condition. This natural variability was intentionally retained to better reflect the demographic structure of field populations. Individuals (males and females) were randomly assigned to experimental units to avoid systematic bias among treatments. There are currently no methods to determine the exact age of adult copepods; therefore, age could not be controlled or included as a factor in the experimental design.

### 2.2 Copper

Copper stock solution (1 g L^-1^) were prepared from CuSO_4_ (purity >99%, Sigma-Aldrich, Søborg, Denmark) following a method similar to that described by Dinh et al. (, 2021, 2020). Copper concentrations in the control treatment (hereafter referred to as the control) were below the detection limit (1 µg L^−1^, ICP-MS).

### 2.3 Experimental design and setup

To test the sensitivity of *P. annandalei* to a range of temperatures and copper exposure, adult copepods were subjected to a fully factorial experimental design comprising 20 treatments: 5 copper concentrations (control, 10, 20, 30, and 40 μg L^-1^) × 4 temperatures (26, 29, 32, and 35°C) (Figure 1). The Cu concentrations varies from ecologically relevant range, covering both low and high levels, as such a range of Cu levels has been found in the coastal waters in Vietnam (up to 63 µg Cu L^−1^, Le and Nguyen, 2018) and worldwide (1.61-560 µg Cu L^−1^, Jonathan et al., 2011, Teasdale et al., 2003). These temperatures were chosen to cover typical temperature ranges in the shallow coastal water in the South of Vietnam, where the study species is abundant (Doan et al., 2018). Each treatment was replicated ten times, resulting in a total of 200 experimental units (50ml Falcon tubes).

**Figure 1.**
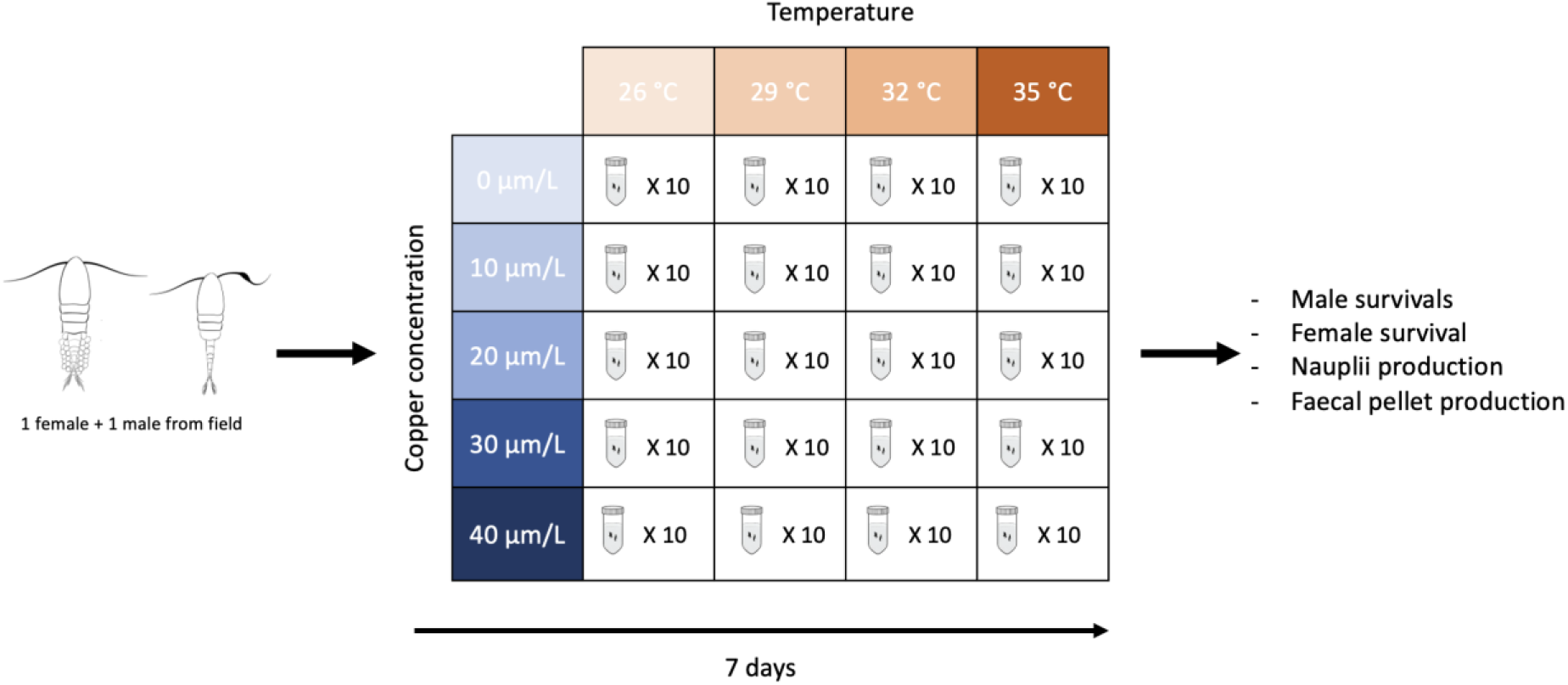
Experimental design used to test the effects of temperature (26, 29, 32, and 35 °C) and copper concentrations (0, 10, 20, 30 and 40 µg L^−1^) on the tropical copepod *Pseudodiaptomus annandalei*.

Each replicate consisted of one egg-carrying female and one male *P. annandalei* placed in a 50 mL Falcon tube containing 20 mL of seawater with the designated copper concentration (equivalent to 100 individuals L^-1^). This density falls within the typical range of 10-400 individuals L^-1^ for natural populations of *P. annandalei* (Truong et al., 2014, Grønning et al., 2019) and is below the density of 400 individuals L^-1^ known to induce density-dependent effects on reproductive performance (Rayner et al., 2017). The tubes were placed in temperature-controlled water baths maintained using aquarium heaters. All copepods were fed with *Isochrysis galbana* at a concentration of 30,000–33,000 cells L^−1^ (equivalent to 800–850 µg C L^−1^; Berggreen et al., 1988; Doan et al., 2018). Throughout the experiment, the salinity, photoperiod, and dissolved oxygen concentration were maintained at 30 PSU which is close to the field conditions at the sampling site (29 PSU; see *Section* 2.1), and a photoperiod was set to 12 h light : 12 h dark. The photoperiod reflects typical tropical conditions in southern Vietnam, where day length remains relatively constant throughout the year. The dissolved oxygen was maintained at approximately 6-7 mg L^-1^, which is close to the saturation levels at experimental temperatures.

To determine the survival of adult *P. annandalei*, cumulative nauplii production, and cumulative faecal pellets, the contents in each bottle were filtered and determined every day (mesh size = 25 μm). Females were not standardized with respect to egg sac developmental stage or clutch size to retain natural variation in reproductive condition and to avoid invasive handling (e.g., detailed microscopic examination or dissection of egg sacs) for repeated measurements. To account for this variability, reproductive output was quantified as cumulative nauplii production over the 7-day experimental period, during which females produced multiple clutches (e.g., the clutch interval ∼29 hours, Golez et al., 2004). The dead copepods were removed from the tube, and their sex was identified. Alive males and females were then returned to the tube with freshly prepared solutions to maintain consistent experimental conditions and to avoid indirect effects of treatments on algal food quality (Holm et al., 2022). The remaining contents, containing nauplii and pellets, were transferred to 12-well plates, fixed with Lugol 4%. Numbers of nauplii and faecal pellets were counted using a stereomicroscope (SZ51, Olympus, Japan). The number of alive females and males, and the total number of nauplii and faecal pellets of each tube after seven days. If one individual died during the experiment, the remaining individual was retained in the same tube, and nauplii production continued to be recorded as females could produce viable nauplii for one or two more clutches (Golez et al., 2004). Faecal pellet production was also quantified and standardised per individual. If both individuals died, reproductive output for that replicate was recorded as zero from that point onward, while cumulative nauplii production from previous days was retained and included in the statistical analyses.

### 2.4 Statistical analyses

Statistical analyses were conducted in R (version 4.4.2). The survival of males and females were analysed using generalized linear mixed models (GLMMs) with a binomial error distribution and logit link function. Temperature, Cu, and their interaction were fitted as fixed effects. Time was initially included in the models as a fixed factor, yet due to limited temporal variations in survival within experimental units (Falcon tubes), it was excluded from the final models. The significance of fixed effects was evaluated using likelihood ratio tests. No significant main or interactive effects were detected; therefore, post hoc comparisons were not conducted for survival.

Cumulative nauplii production and cumulative faecal pellets were analysed using generalized linear models (GLMs) with a negative binomial error distribution as their counts showed overdispersion under a Poisson model (dispersion ∼15). A log link function to ensure positive fitted values appropriate for cumulative count data (Truong et al., 2022). Temperature, Cu, and their interaction were fixed factors. Model significance was assessed using Type III Wald Chi-square tests. Post hoc comparisons were conducted only when significant main effects (temperature or copper) or interaction effects were detected using Tukey’s HSD test to identify differences among treatments while controlling Type I error rates. To evaluate temporal dynamics of daily nauplii and faecal pellet production during the experimental period, two separate GLMs were conducted with the same error structure as described above, with Time included as an additional fixed factor along with its interactions with Temperature and Cu (see Appendix S2). When significant temporal interactions were detected, selected pairwise comparisons were conducted using Tukey’s HSD test to aid interpretation of treatment differences at specific time points and to support data visualisations.

In all models, individual-level factors such as age and physiological condition were not included as explanatory variables. Instead, natural variability among individuals was intentionally retained to better reflect the demographic structure of field populations and to capture variability in sensitivity to environmental stressors.

## 3. Results and discussion

### 3.1 Effects of temperature on P. annandalei

Survival did not differ among all 20 temperature × Cu treatment combinations during this short-term exposure (Table 1, Figures 2 and 3). In our previous studies, temperatures of 34 - 35 °C caused reductions in survival when the exposure duration lasted for an entire one or two generations (20-24 days) of *P. incisus* a congeneric species (Dinh et al., 2021), suggesting a species specific vulnerability and also the short-term tolerance. In a short-term exposure, copepods may prioritize maintenance to preserve survival under acute heat stress (Sokolova, 2013). Such prioritization often comes at energetic costs, which are often manifested in energy and resources limitation (Metcalfe, 2025, Sokolova, 2013, Saiz et al., 2022). The reduction in feeding activity at 35 °C, as indicated by decreased faecal pellet production suggests that energy intake may not fully match the elevated metabolic demand associated with warming (Metcalfe, 2025, Sokolova, 2013, Saiz et al., 2022). Faecal pellet production is widely used as a proxy for feeding activity in copepods and is generally correlated with ingestion rates, but it also depends on gut passage time and assimilation efficiency, both of which can be influenced by temperature and contaminant exposure (Besiktepe and Dam, 2002, Dinh et al., 2019, Doan et al., 2018). As a result, changes in pellet production represent integrated responses of feeding and digestive processes (e.g., Besiktepe and Dam, 2002, Dinh et al., 2019, Doan et al., 2018). Under heat stress, available energy may be allocated to more essential maintenance processes, such as the upregulation of heat shock proteins, and coping with oxidative damage (Harada et al., 2019, Low et al., 2018), potentially trading off against energetically expensive processes such as reproduction (e.g., Siegle et al., 2022, Ginther et al., 2024).

**Table 1.** The results of the statistical analyses testing effects of heat stress (Temp) and copper concentration (Cu) on survival of males and females, cumulative nauplii, and cumulative fecal pellets of *Pseudodiaptomus annandalei*. Significant *P* values are signed in bold.

| Effects | Male survival |  |  | Female survival |  |  | Cumulative nauplii<br>production |  |  | Cumulative faecal pellet<br>production |  |  |
| --- | --- | --- | --- | --- | --- | --- | --- | --- | --- | --- | --- | --- |
| | df1, df2 | F | *P* | df1, df2 | *F* | *P* | df | $\chi^2$ | *P* | df | $\chi^2$ | *P* |
| Temp | 3, 180 | 2.493 | 0.075 | 3, 180 | 0.226 | 0.878 | 3 | 11.806 | **0.008** | 3 | 13.679 | **0.003** |
| Cu | 4, 180 | 0.389 | 0.610 | 4, 180 | 1.637 | 0.167 | 4 | 6.153 | 0.188 | 4 | 3.794 | 0.434 |
| Temp × Cu | 12, 180 | 1.366 | 0.184 | 12, 180 | 1.254 | 0.249 | 12 | 29.650 | **0.003** | 12 | 47.788 | **<0.001** |

**Figure 2.**
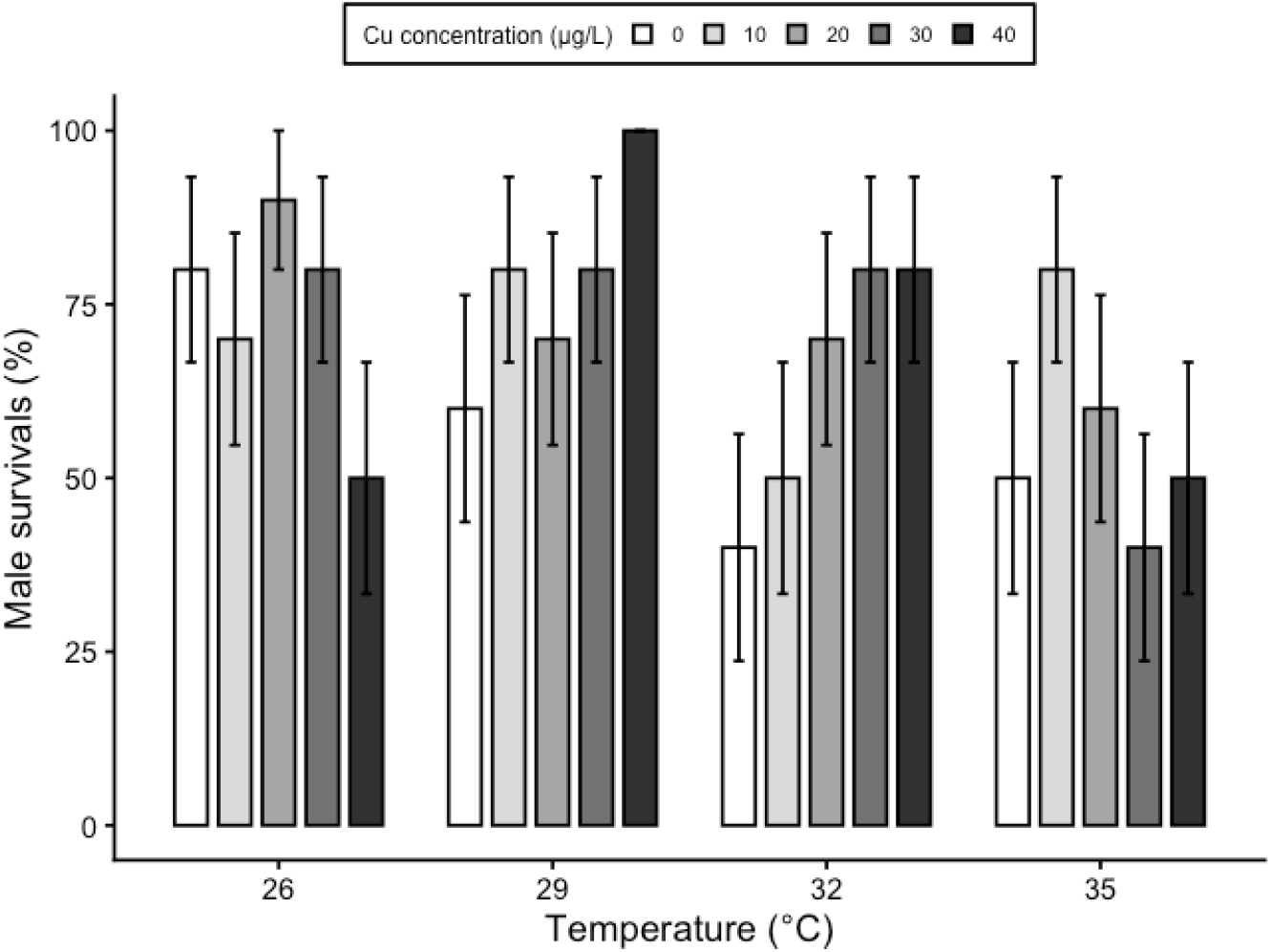
Effects of temperature and copper on male survival of *Pseudodiaptomus annandalei*. Data are visualised as mean ± SEs (n = 10 replications).

**Figure 3.**
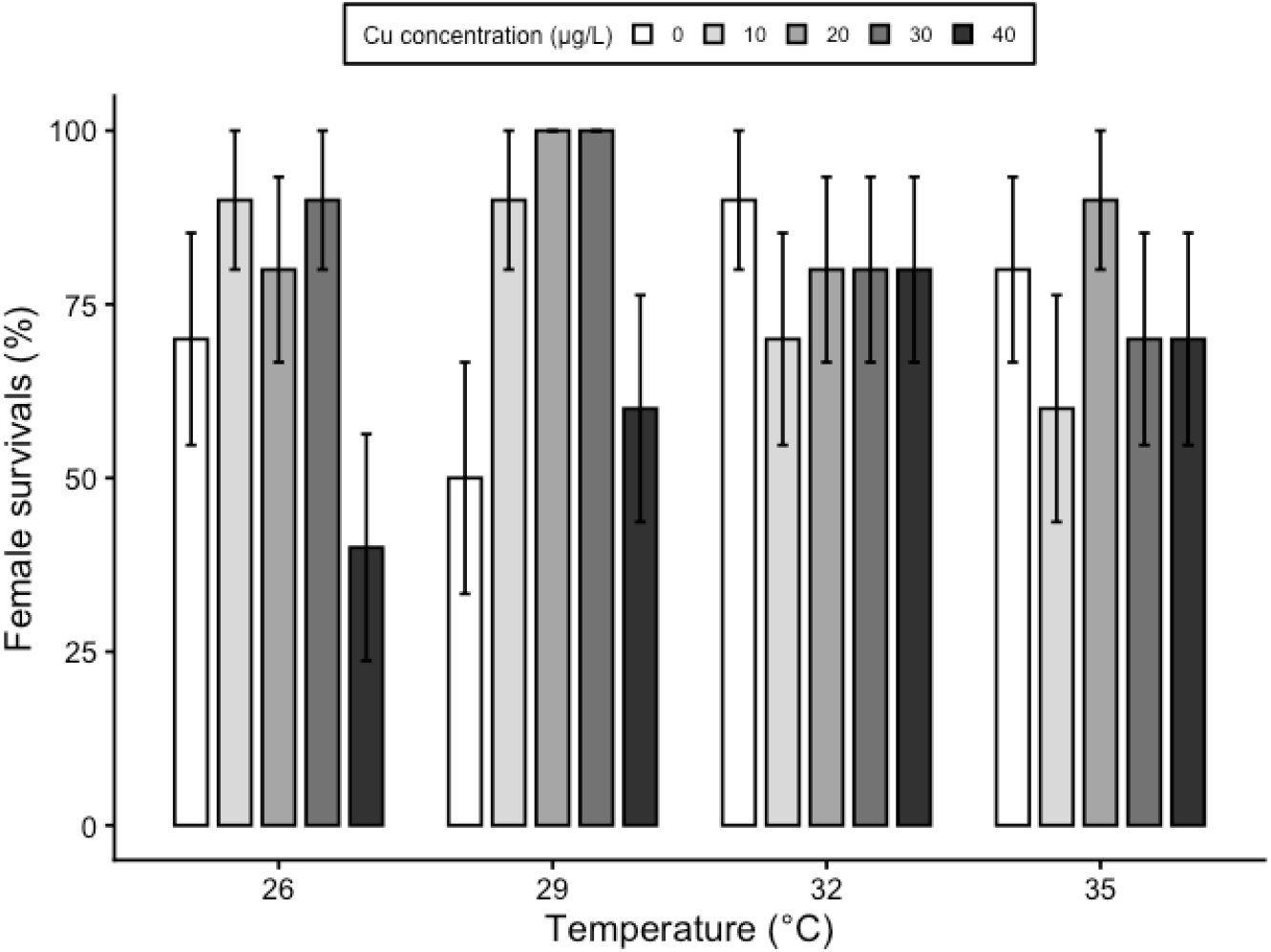
Effects of temperature and copper on female survival of *Pseudodiaptomus annandalei*. Data are visualised as mean ± SEs (n = 10 replications).

Heat stress significantly reduced the cumulative nauplii and faecal pellet production of *P. annandalei*. Highest cumulative nauplii produced by *P. annandalei* was found at 32 °C, which is approximately 11% higher than those at 29 °C, and 26% higher than the lowest ones at 26 °C and 35 °C (main temperature effect, Table 1, Figure 4). Although daily nauplii production did not differ consistently among temperatures as indicated by no main effect of temperature on the daily nauplii production (see Table S2.1, Figure S2.1 and Appendix S2), these differences emerged cumulatively over time, reflecting the integration of small, time-dependent variations in reproductive output. These cumulative differences suggest that 32 °C supports more sustained reproductive performance during later stages of the production period (Figure S2.1). In contrast, extreme temperature (35 °C) likely imposes increasing cumulative physiological stress during prolonged exposure, reducing reproductive output over successive days and resulting in lower cumulative production.

**Figure 4.**
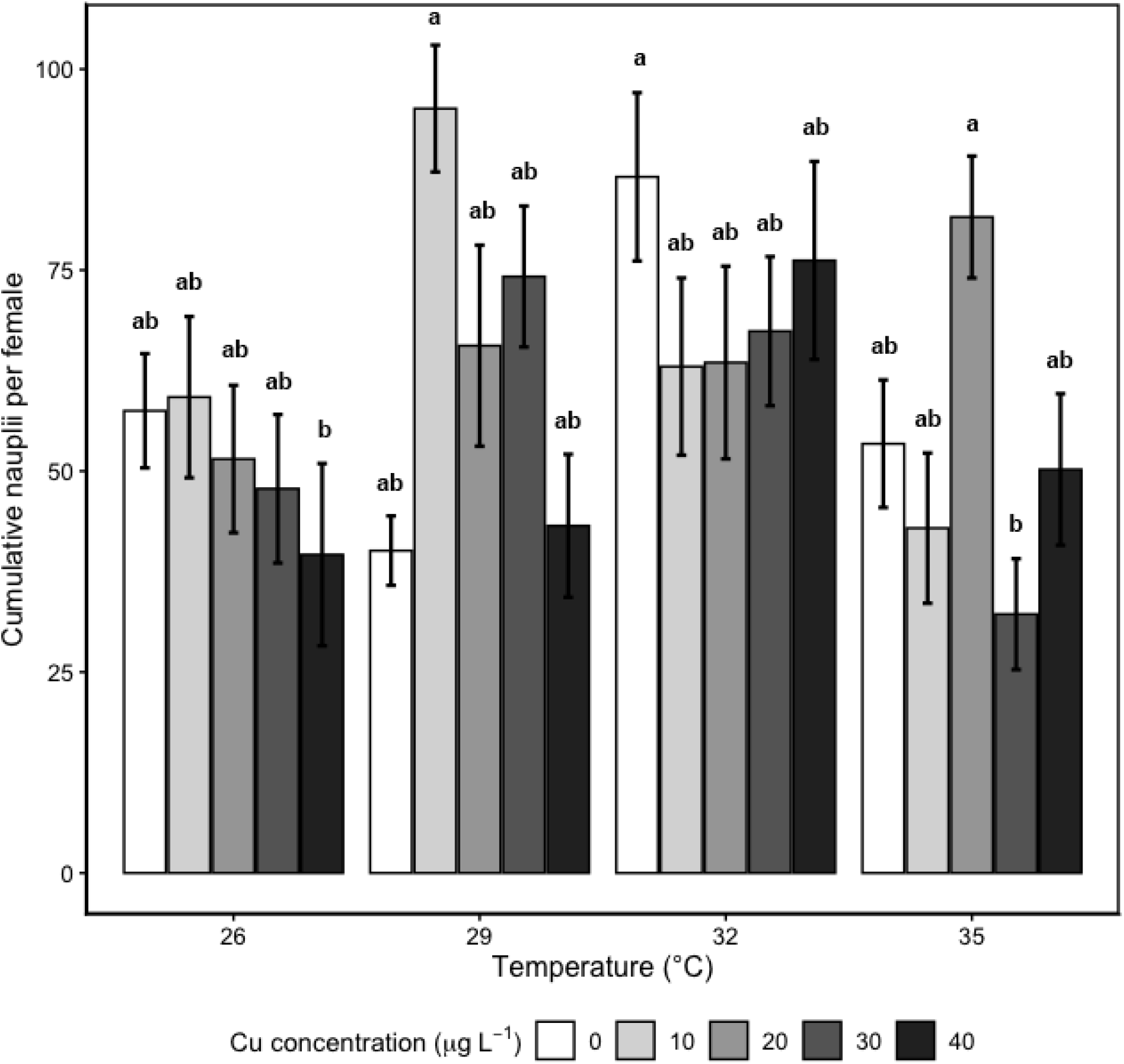
Effects of temperature and copper on cumulative nauplii production per female of *Pseudodiaptomus annandalei*. Data are visualised as mean ± SEs (n = 10 replications). Different letters denote statistically significant differences (Tukey HSD, *P* < 0.05).

Across all Cu concentrations, the cumulative number of pellets per individual was highest at 32 °C, approximately 10% higher than at 35 °C and 16% higher than those at 26 °C and 29 °C (main temperature effect, Table 1, Figure 5). The unimodal patterns in cumulative nauplii and faecal pellet production partially support H1 as it was primarily evident when responses were averaged across Cu concentrations (see further details in section 3.2). These results are consistent with previous findings that 32 °C represents the upper optimal thermal range of for *P. annandalei* (Lehette et al., 2016). Temporal analyses further showed that feeding rates were highest at 32 °C during the later stages of the experiment, indicating that temperature effects on feeding developed progressively. Specifically, feeding rates increased at 32 °C, remained relatively constant at lower temperatures, and were constrained at 35 °C on days 6 and 7. At lower temperatures, feeding was likely limited by higher attacking and handling time due to higher viscosity and slower gut throughput and longer processing times (Tyrell and Fisher, 2019, Dam and Peterson, 1988), whereas at 35 °C, increased physiological stress may have constrained feeding (Saiz et al., 2022). As faecal pellet production scales with ingestion and carbon assimilation (Besiktepe and Dam, 2002), suggesting a higher energy acquisition at 32 °C than at both at lower or higher temperatures. Temperature of 35 °C exceeds the upper thermal optimum of this species (Lehette et al., 2016), which is in agreement with general findings of the reduced feeding in tropical copepods of the genus *Pseudodiaptomus* (Doan et al., 2018, 2019, Nguyen et al., 2020a, 2020b). This could create a double energetic constraint: increased expenditure combined with reduced intake efficiency, that may partly contribute to reduced reproduction (Sokolova and Lannig, 2008, Holmstrup et al., 2010, Jackson et al., 2016, Dinh et al., 2022). Such temperature-dependent constraints on feeding and reproduction may have broader ecological implications, potentially reducing population productivity and altering the trophic role of copepods in coastal planktonic food webs (Doan et al., 2019, Chew et al., 2012).

**Figure 5.**
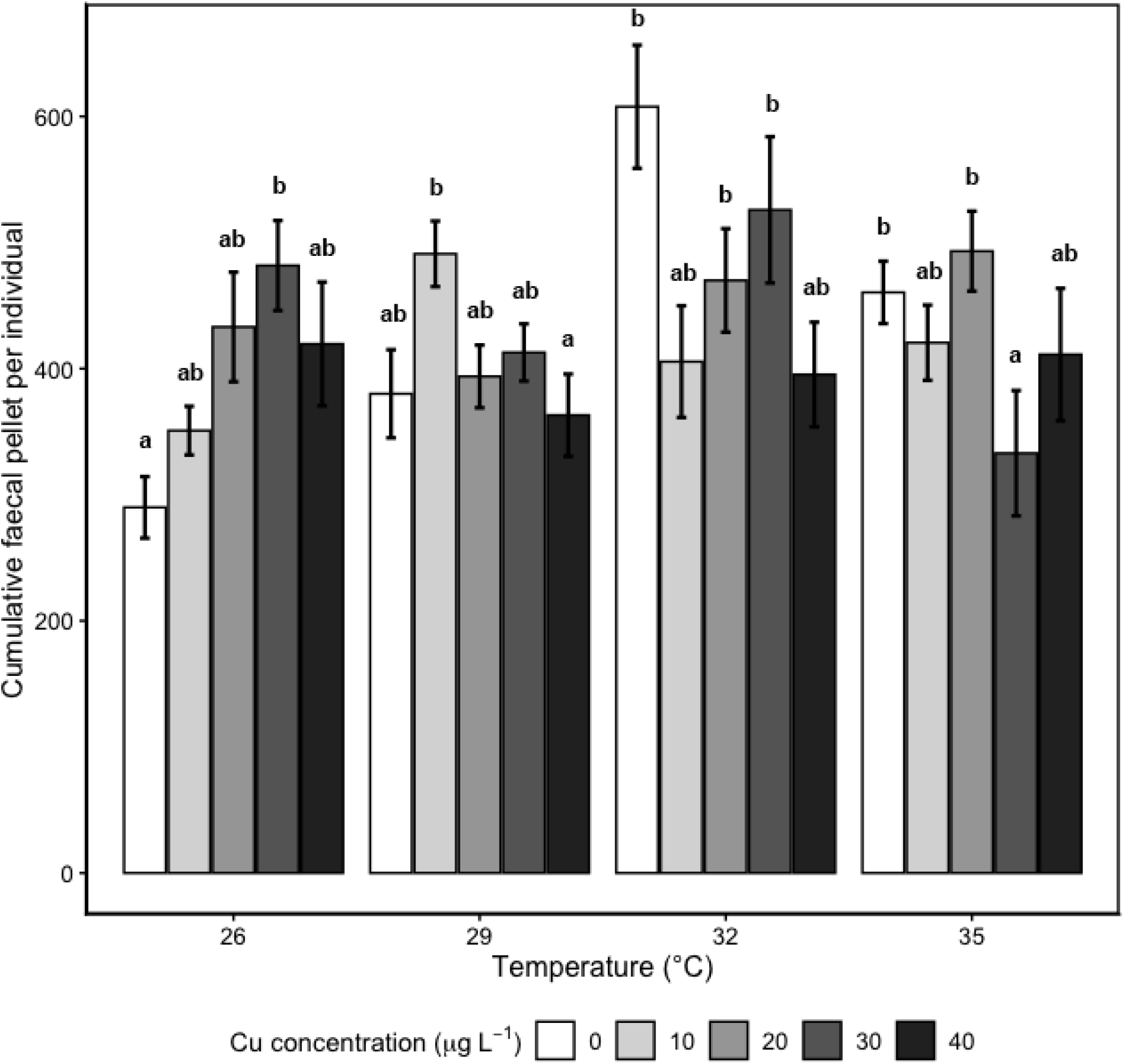
Effects of temperature and copper on cumulative fecal pellet production per individual of *Pseudodiaptomus annandalei*. Data are visualised as mean ± SEs (n = 10 replications). Different letters denote statistically significant differences (Tukey HSD, *P* < 0.05).

### 3.2 Temperature dependent Cu-toxicity in P. annandalei

Contrary to H2, Cu exposure alone did not significantly affect the survival of male or female *P. annandalei*, nor cumulative nauplii and faecal pellet production during the seven-day experiment (Table 1, Figures 2-5). Survival varied considerably among temperature × Cu treatments, but no consistent pattern was observed across treatments, that might be associated with high variabilities among replicates (Figures 2 and 3). Such patterns likely reflect differences in reproductive stage, and physiological condition (Saiz et al., 2015, Truong et al., 2020), which are inherent to natural populations. This variability may reduce detectability of potential temperature and Cu effects on survival, which has been found in our previous studies on *P. annandalei* and *P. incisus* at comparable Cu concentrations (Dinh et al., 2021, Dinh et al., 2020).

High variabilities of treatment-specific responses of *P. annandalei* were observed for cumulative nauplii and faecal pellet production, that may be explained by cumulative heat stress and physiological specific performance. At 32 °C, cumulative responses did not differ significantly among Cu concentrations (Tukey HSD, *P* > 0.05), whereas significant variability was observed at both lower and higher temperatures. At 35 °C cumulative nauplii production was only lower at 30 µg Cu L^−1^, and highest at 20 µg Cu L^−1^, with intermediate values were observed at other concentrations (Temperature × Cu interaction, Table 1, Figure 4). For faecal pellet production, *P. annandalei* showed significantly higher value in non-Cu exposed copepods at 32 °C and lower values in non-Cu exposed copepods at 26 °C, as well as in copepods exposed to 40 µg Cu L^−1^ at 29 °C and 30 µg Cu L^−1^ at 35 °C (Temperature × Cu interaction, Table 1, Figure 5). Intermediate values were observed in other conditions and were not significantly different from either the highest or lowest treatments. These results only partially support H3.

Temporal analyses provided a more detailed of time-dependent treatment effects over the experimental period. Daily nauplii production per female showed significant interactions with temperature and Cu over time (significant Temperature × Time and Temperature × Time × Cu interactions; Table S1, Appendix S2). Specifically, nauplii production did not differ among treatments during days 1–5, indicating limited short-term effects of temperature and Cu on reproductive output (Table S2.1, Figure S2.1). Differences emerged during the later stages (days 6–7), but in an inconsistent manner. At 29 °C, nauplii production was highest in copepods exposed to 10 µg Cu L^−1^ and lowest in non-Cu-exposed individuals on days 6-7 (Figure S2.1). At 32 °C, nauplii production was generally higher in non-Cu-exposed copepods than in those exposed to Cu (10–40 µg Cu L^−1^) on day 6. At 35 °C, nauplii production remained low across all treatments. These inconsistent time- and Cudependent responses, particularly at 35 °C, suggest that reproductive performance was not only reduced but also highly variable under supra-optimal temperature conditions. These patterns imply that the combined effects of temperature and Cu were context-dependent and became apparent only after prolonged exposure, challenging the ecological risk assessment of pollutants under climate change (Moe et al., 2013, Dinh et al., 2022, Sokolova and Lannig, 2008).

Variability in male survival may have contributed to variation in reproductive output among treatments as the loss of males may reduce mating success and consequently limit nauplii production (Kiørboe, 2007, Gusmao and McKinnon, 2009). However, in the present study, survival of both males and females did not differ among temperature and Cu treatments, indicating that mortality did not vary systematically across treatments and therefore did not bias comparisons of reproductive output. Reproductive output was quantified cumulatively over the experimental period, such that outcomes inherently reflect the combined effects of survival and reproductive performance within each replicate.

Faecal pellet production per individual showed significant temporal variation and interactions with both temperature and Cu (main effect of Time, and significant Temperature × Time, Time × Cu, and Temperature × Time × Cu interactions; Table 1, Appendix S2). Specifically, non-Cu-exposed copepods showed lower faecal pellet production than Cu-exposed individuals only on day 5 at 26 °C and day 7 at 29 °C, but higher faecal pellet production on day 5 at 29 °C (Table S2.1, Figure S2.2). This may reflect a late-stage increase in feeding under Cu exposure at these temperatures. In contrast, different temporal patterns were observed at higher temperatures. At 32 °C, non-Cu-exposed copepods showed higher faecal pellet production on day 1, and this pattern re-emerged during the later stages of the experiment (day 6) (Table S2.1, Figure S2.2). At 35 °C, copepods exposed to 10, 20 and 40 µg Cu L^−1^ produced more faecal pellets than those in 0 and 30 µg Cu L^−1^ on day 1 (Tukey HSD, *P* < 0.05, but these differences were not sustained over subsequent days (days 2–7) (Table S2.1, Figure S2.2). Together, these results indicate that feeding responses were both temperature- and time-dependent, with early responses at higher temperatures being transient, while delayed responses diverged across temperature conditions. Specifically, sustained increases in feeding were observed near the thermal optimum (32 °C), whereas responses at supra-optimal temperature (35 °C) were more constrained and not maintained over time.

Importantly, our results also suggest that Cu did not shift the thermal optimum but influenced performance in a time- and condition-dependent manner, but inconsistently to a dose-dependent manner, particularly under extreme temperatures beyond the thermal optimum. This pattern is consistent with recent ecotoxicological studies showing that contaminant effects may become more variable or pronounced toward thermal extremes, although such effects are not always uniform across treatments (Dinh and Vu, 2025). In tropical systems, where organisms already live close to their upper thermal limits (Deutsch et al., 2008, Lehette et al., 2016), even small temperature increases may amplify contaminant effects. In the present study, sublethal endpoints such as feeding and reproductive output showed greater variability across treatments than survival, indicating that these traits may respond more sensitively to stressors, although consistent Cu effects were limited.

### 3.3 Implications for ecological risk assessments and conclusions

Our findings have several implications for understanding and managing the impacts of multiple environmental stressors in tropical coastal ecosystems. In particular, they provide empirical support for the growing recognition that contaminant effects cannot be evaluated independently of environmental temperature, a central issue in ecotoxicology under global change (Moe et al., 2013, Sokolova and Lannig, 2008, Holmstrup et al., 2010, Dinh et al., 2022, Noyes et al., 2025). The influence of Cu on reproductive output became evident under sup-optimal thermal conditions, suggesting that contaminant toxicity may only emerge under particular environmental contexts, such as during marine heatwaves or other extreme events. Consequently, ecological risk assessments based on single-temperature toxicity tests may underestimate the vulnerability of tropical marine organisms under future warming scenarios.

While our experiment was conducted under constant temperature conditions, natural environments are characterised by temporal variability, including diel fluctuations (Doan et al., 2018) and marine heatwaves. Such variability may further influence organismal responses and modify stressor interactions. Our results therefore provide a baseline for understanding how temperature-dependent contaminant effects may manifest under more complex and dynamic thermal regimes and highlight the need for future studies incorporating realistic temperature variability. The copepods of the genus *Pseudodiaptomus* play a central role in tropical coastal food webs by transferring primary production from phytoplankton to higher trophic levels, including fish larvae and juvenile stages (Chew et al., 2012). Changes in feeding activity and reproductive output can therefore propagate through the food web and influence energy transfer in coastal ecosystems (Chew et al., 2012, Turner, 2004).

Finally, the relatively high variability observed among treatments likely reflects the natural heterogeneity of wild populations (Blanda et al., 2015, Grønning et al., 2019). Field-collected copepods may differ in age, reproductive stage, and physiological condition, all of which can influence their responses to environmental stressors (Muyssen and Janssen, 2007, Dinh et al., 2022, Schmid et al., 2023, Rohr et al., 2016). While such variability can obscure treatment effects in short-term laboratory experiments, it may better represent an inherent and ecological meaningful feature of natural populations. In this context, our results suggest that environmentally realistic population variability may mask stressor effects that could become more pronounced under extreme environmental conditions, further challenging the ecological risk assessments. Therefore, combining controlled laboratory studies with standardised life stages and physiological conditions, with ecologically realistic experiments, and field observations will be necessary to improve predictions of how multiple stressors influence tropical marine ecosystems under ongoing global change (Dinh et al., 2022).

## Ethics declarations

This manuscript has not been published previously. This manuscript is not under consideration for publication elsewhere. The article’s publication is approved by all authors. If accepted, the article will not be published elsewhere in the same form, in English, or any other language, including electronically, without the written consent of the copyright holder.

## Acknowledgments

We thank Nha Trang University for supporting facilities to conduct this study and to implement the project B2024-TSN-15.

