## Appendix for "Feeding and reproduction of a tropical coastal copepod across warming and copper gradients": Appendix S2.docx

**Table S2.1**. The results of the statistical analyses testing effects of heat stress (Temp) and copper concentration (Cu) on daily nauplii produced per female, and daily fecal pellets per individual of *Pseudodiaptomus annandalei*. Significant *P* values are signed in bold.

| Effects | Daily nauplii production | | | Daily faecal pellet production | | | |
| --- | --- | --- | --- | --- | --- | --- | --- |
|  | df | χ² | *P* | df | χ² | *P* | |
| Temp | 3 | 3.939 | 0.268 | 3 | 13.694 | | **0.003** |
| Cu | 4 | 0.891 | 0.926 | 4 | 0.805 | | 0.938 |
| Temp × Cu | 12 | 7.584 | 0.817 | 12 | 24.482 | | **0.017** |
| Time | 6 | 10.100 | 0.121 | 6 | 14.751 | | **0.022** |
| Time × Temp | 18 | 36.64 | **0.006** | 18 | 36.676 | | **0.006** |
| Time × Cu | 12 | 7.584 | 0.817 | 24 | 43.046 | | **0.009** |
| Time × Temp × Cu | 72 | 109.964 | **0.003** | 72 | 125.045 | | **<0.001** |

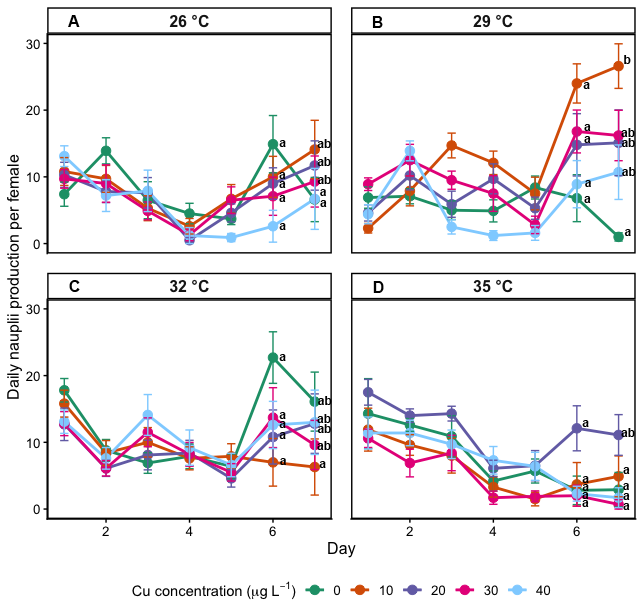

**Figure S2.1**. Daily nauplii production per female of Pseudodiaptomus annandalei across temperatures (26, 29, 32, and 35 °C) and Cu concentrations (0, 10, 20, 30, and 40 µg L⁻¹) over the 7-day exposure period. Different letters indicate statistically significant differences among treatments on a given day (Tukey’s HSD, P < 0.05). The absence of letters indicates no significant differences on that day.

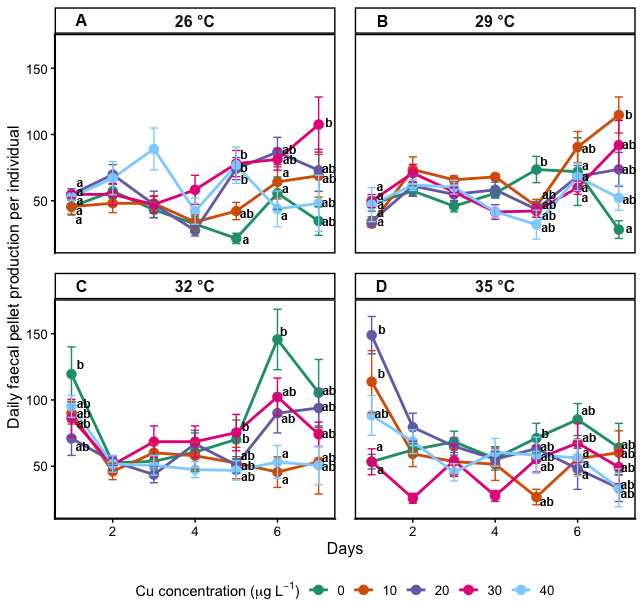

**Figure S2.2**. Daily faecal pellet production per individual of Pseudodiaptomus annandalei across temperatures (26, 29, 32, and 35 °C) and Cu concentrations (0, 10, 20, 30, and 40 µg L⁻¹) over the 7-day exposure period. Different letters indicate statistically significant differences among treatments on a given day (Tukey’s HSD, P < 0.05). The absence of letters indicates no significant differences on that day.
